# Discovery of viral myosin genes with complex evolutionary history within plankton

**DOI:** 10.1101/2021.03.14.435220

**Authors:** Soichiro Kijima, Tom O. Delmont, Urara Miyazaki, Morgan Gaia, Hisashi Endo, Hiroyuki Ogata

## Abstract

Nucleocytoplasmic large DNA viruses (NCLDVs) infect diverse eukaryotes and form a group of viruses with capsids encapsulating large genomes. Recent studies are increasingly revealing a spectacular array of functions encoded in their genomes, including genes for energy metabolisms, nutrient uptake, as well as cytoskeleton. Here, we report the discovery of genes homologous to myosins, the major eukaryotic motor proteins previously unrecognized in the virosphere, in environmental genomes of NCLDVs from the surface of the oceans. Interestingly, these genes were often accompanied by kinesin genes in the environmental genomes, suggesting a role of these viral proteins in the intracellular viral particle transport. Phylogenetic analyses indicate that most viral myosins (named “virmyosins”) belong to the *Imitervirales* order, except for one belonging to the *Phycodnaviridae* family. On the one hand, the phylogenetic positions of virmyosin-encoding *Imitervirales* are scattered within the *Imitervirales*. On the other hand, *Imitervirales* virmyosin genes form a monophyletic group in the phylogeny of diverse myosin sequences. Furthermore, phylogenetic trends for the virmyosin genes and viruses containing them were incongruent. Based on these results, we argue that multiple transfers of myosin homologs have occurred not only from eukaryotes to viruses but also between viruses, supposedly during co-infections of the same host.

## INTRODUCTION

Viruses were considered as tiny and simple biological objects until La Scola et al. discovered a giant virus from the water of a cooling tower in 2003 (Scola et al., 2003). The virus named mimivirus is 750 nm in particle size and possesses a 1,182 kbp genome (Colson et al., 2017), a dimension that was large and complex enough to blow off the classical perception of viruses. After the discovery of mimivirus, related viruses were isolated including marseilleviruses, pandoraviruses, and pithoviruses, many of them with similar or even larger-sized particles or genomes (Abergel et al., 2015; Colson et al., 2017). These giant viruses infect diverse eukaryotes, possess a double-stranded DNA genome, belong to the phylum *Nucleocytoviricota* (Koonin et al., 2020), and are commonly referred to as nucleocytoplasmic large DNA viruses (NCDLVs) (Iyer et al., 2006). The monophyletic origin of NCLDVs has been suggested based on the presence of about 40 core genes of NCLDVs that can be traced back to their putative last common ancestor (Koonin and Yutin, 2019) as well as the congruent phylogenies of the most conserved 8 proteins responsible for virion morphogenesis and informational processes (Guglielmini et al., 2019).

Because of their large virions, NCLDVs can encapsulate a large genome (several hundred kb up to 2.5 Mb) in their particles. Smaller viruses (such as small RNA viruses) encode only genes that are essential for their genome replication and capsid formation, whilst NCDLVs encode numerous genes that are not directly involved in their genome replication and virion morphogenesis (Moniruzzaman et al., 2020). These genes, often called auxiliary metabolic genes, are considered to function in reprogramming host metabolism and molecular machinery during viral infection towards enhancing viral replication and subsequent transmission to another host. For example, the recently characterized Prymnesium kappa virus RF01 encodes genes for all four succinate dehydrogenase subunits, as well as genes for modulating β-oxidation pathway (Blanc-Mathieu et al., 2021). These viral genes are suggested to boost energy production during viral replication, which can deteriorate host metabolism, or to enhance the supply of building blocks for viral replication. Another recent study reported the presence of actin genes (viractins) in NCLDV genomes (Cunha et al., 2020). Viractins are hypothesized to help viral infections by controlling the localization of the viral factory close to the host nucleus.

Hundreds of genomes have already been sequenced for cultured NCLDVs, yet these represent only the tip of iceberg of the diverse NCLDVs uncovered through environmental surveys (Schulz et al., 2020). To by-pass cultivation, genome-resolved metagenomics has been applied to large metagenomic surveys, including on oceanic samples collected by *Tara* Oceans (Sunagawa et al., 2020), in order to characterize NCLDV metagenome-assembled genomes (MAGs) containing the gene pool of thousands of those viruses (Moniruzzaman et al., 2020; Schulz et al., 2020). NCLDV MAGs revealed a cosmopolitan nature of these viruses, extensive gene transfers with eukaryotes, and their complex metabolic capabilities.

In this study, we describe the identification of myosin genes in previously published NCLDV MAGs as well as newly identified ones derived from a manual binning and curation effort focused on large cellular size fractions of *Tara* Oceans enriched in NCLDVs when infecting planktonic eukaryotes (see Materials and Methods). Myosin genes, which have not been previously described in viral genomes from cultures, form a superfamily of motor proteins involved in a wide range of motility processes in eukaryotic cells. Myosins have been grouped into various classes (Odronitz and Kollmar, 2007). Most myosins are classified into class 1 to 35 based on their phylogenetic relationships (Odronitz and Kollmar, 2007), whilst other myosins are phylogenetically orphan and classified into class A to U (Odronitz and Kollmar, 2007). The functions of orphan myosins are often unknown, while the functions of some members of the class 1 to 35 are characterized. For example, some myosins of class 2 contract muscle (Pertici et al., 2018), whilst some myosins of class 5 transport specific material along actin filaments (Hammer and Sellers, 2012). The various functions of myosins are supported by the head domains, which are universally conserved among myosins, interact with actins, and frequently serve for phylogenetic analyses. The head domains of myosins interact with actins when myosins bind ADP, whereas, when ADP is absent or ATP is in the ADP-binding site of myosins, the head domains no longer interact with actins.

## MATERIALS AND METHODS

### NCLDV MAGs derived from the *Tara* Ocean Project

Newly identified NCLDV metagenome-assembled genomes (MAGs) were manually characterized and curated from the *Tara* Oceans metagenomes (size fractions > 0.8 µm), based on an initial binning strategy at large-scale focused on eukaryotes (Delmont et al., 2020), and following the same workflow as in previous studies (Cunha et al., 2020; Kaneko et al., 2020; Delmont et al., 2021). Briefly, metagenomes were organized into 11 sets based on their geography, and each set was co-assembled using MEGAHIT (Li et al., 2015) v.1.1.1. For each set, scaffolds longer than 2.5 kbp were processed within the bioinformatics platform anvi’o v.6.1 (Eren et al., 2015) to generate genome-resolved metagenomes (Delmont et al., 2018). CONCOCT (Alneberg et al., 2014) was used to identify large clusters of contigs within the set. We then used HMMER (Eddy, 2011) v3.1b2 to search for eight NCLDV gene markers (Guglielmini et al., 2019), and identified NCLDV MAGs by manually binning CONCOCT clusters using the anvi’o interactive interface. Finally, NCLDV MAGs were manually curated using the same interface, to minimize contamination as described previously (Delmont and Eren, 2016).

### Sequence datasets

To prepare a sequence set for the DNA polymerase elongation subunit family B (PolB), we first extracted PolB sequences from NCVOG (Yutin et al., 2009). Next, we collected PolB sequences by performing BLASTP from Virus-Host DB (Mihara et al., 2016) against the PolB sequences from NCVOG. We retained hits with an E-value < 1e-10. To identify PolBs in the NCLDV MAGs, we performed BALSTP from sequences derived from NCDLV MAGs generated by Moniruzzaman et al. (Moniruzzaman et al., 2020), Schulz et al. (Schulz et al., 2020) and ourselves (*Tara* Oceans MAGs) against the PolB sequences from NCVOG and Virus-Host DB. We retained hits with an E-value < 1e-10 and with their length in a range from 800 amino acids (aa) up to 1800 aa. We pooled these PolB sequences, and then removed redundancy using cd-hit (4.8.1) (Li and Godzik, 2006) and manually curated the dataset to reduce its size.

For myosin homologs, we used full-length myosin sequences from a previous study (Odronitz and Kollmar, 2007) as primal references for myosins. This dataset contains various classes of myosins from diverse organisms. Next, we extracted myosin homologs from MMETSP (Keeling et al., 2014) and RefSeq (O’Leary et al., 2016) by performing BLASTP (blast 2.11.0) against the primal references to generate secondary references for full-length myosin sequences. We considered hits with an E-value < 1e-10 in this search. We identified myosin head domains in the primal and secondary myosin reference sequences by performing hmmscan (HMMER 3.3.1) (Eddy, 2011). We used Pfam (El-Gebali et al., 2019) as the HMM model for the hmmscan search and considered hits that were annotated as “Myosin_head” and showed an E-value < 1e-10. We filtered out the primal and secondary reference sequences to reduce redundant sequences. To identify myosin sequences in the NCLDV MAGs, we performed BLASTP from sequences derived from MAGs against the primal myosin references. We considered hits with an E-value < 1e-10. We identified myosin head domains by performing hmmscan (Eddy, 2011). We considered hits that were annotated as “Myosin_head” with an E-value < 1e-10 and length longer than 550 aa and shorter than 800 aa.

### Multiple sequence alignment

Multiple sequence alignments were generated using MAFFT (v7.471) (Katoh et al., 2019) with the L-INS-i algorithm, which is suitable for sequences that have only one alignable domain. After multiple alignment, we removed gappy sites by using trimal (v1.4.rev15) (Capella-Gutiérrez et al., 2009) with “-gappyout” option.

For myosin sequences from viruses and eukaryotes, we generated two trimmed alignments by removing gapped columns using “-strict” option (named “strict dataset”), in addition to the “-gappyout” option (named “gappy dataset”). Finally, we removed sequences which have more than 30% gaps along the entire length of the alignment from each dataset.

### Phylogenetic analysis

The phylogenetic analyses were conducted with the ML framework using IQ-TREE (1.6.12) (Minh et al., 2020). For each alignment, we used the best substitution model selected by IQ-TREE. The selected models are described in the figure legends. The branch support values were computed based on non-parametric bootstrap method with 100 bootstrap replicates.

For myosin sequences from viruses and eukaryotes, we also built phylogenetic trees using RAxML (8.2.12) (Stamatakis, 2014) on both “gappy” and “strict” datasets. Therefore, we obtained four trees for these myosin sequences (i.e., gappy/IQ-TREE, strict/IQ-TREE, gappy/RAxML, and strict/RAxML).

### Visualization

The phylogenetic trees were visualized with iTOL v4 (Letunic and Bork, 2019).

### Statistical test for co-existing genes

We performed InterProScan (5.44-79.0) (Mitchell et al., 2019) to find motifs against each protein sequence derived from the NCLDV MAGs. We counted the number of each motif discovered in *Imitervirales*, which was annotated based on the PolB phylogenetic tree. Next, we conducted chi-square test with the Benjamini-Hochberg correction to set the false discovery rate at 0.01 to define the motifs that were significantly enriched in the *Imitervirales* encoding virmyosins compared with other *Imitervirales*.

## RESULTS AND DISCUSSION

### Myosin genes in NCLDV genomes

We identified myosin-related genes in a total of 24 NCLDV MAGs (out of 2,275 considered in our survey) by performing BLASTP against reference myosin sequences compiled from a previous study (Odronitz and Kollmar, 2007), through our own effort independent from a parallel environmental genomic survey (Ha et al., 2021). These NCLDV MAGs were derived from the large cellular size fractions of *Tara* Oceans (n=10) and studies by Moniruzzaman et al. (n=5) and Schulz et al. (n=9) (Moniruzzaman et al., 2020; Schulz et al., 2020). All NCLDV MAGs but one originate from marine plankton samples (**Table S1**). The viral sequences contain the head domain but do not show the tail domain. An alignment of myosin head domain sequences from various organisms with these viral sequences establishes their strong homology (**Figure S1**). The taxonomies of the closest homologs of other genes encoded together with the myosin homologs in the same contigs (**Figure 1**) revealed that a large proportion of genes (up to 44% and 15% on average) in the contigs best match to NCLDV genes, thus excluding the possibility of contaminated myosin homologs in the NCLDV MAGs. We designate the viral myosin homologs as virmyosins.

**Figure 1.**
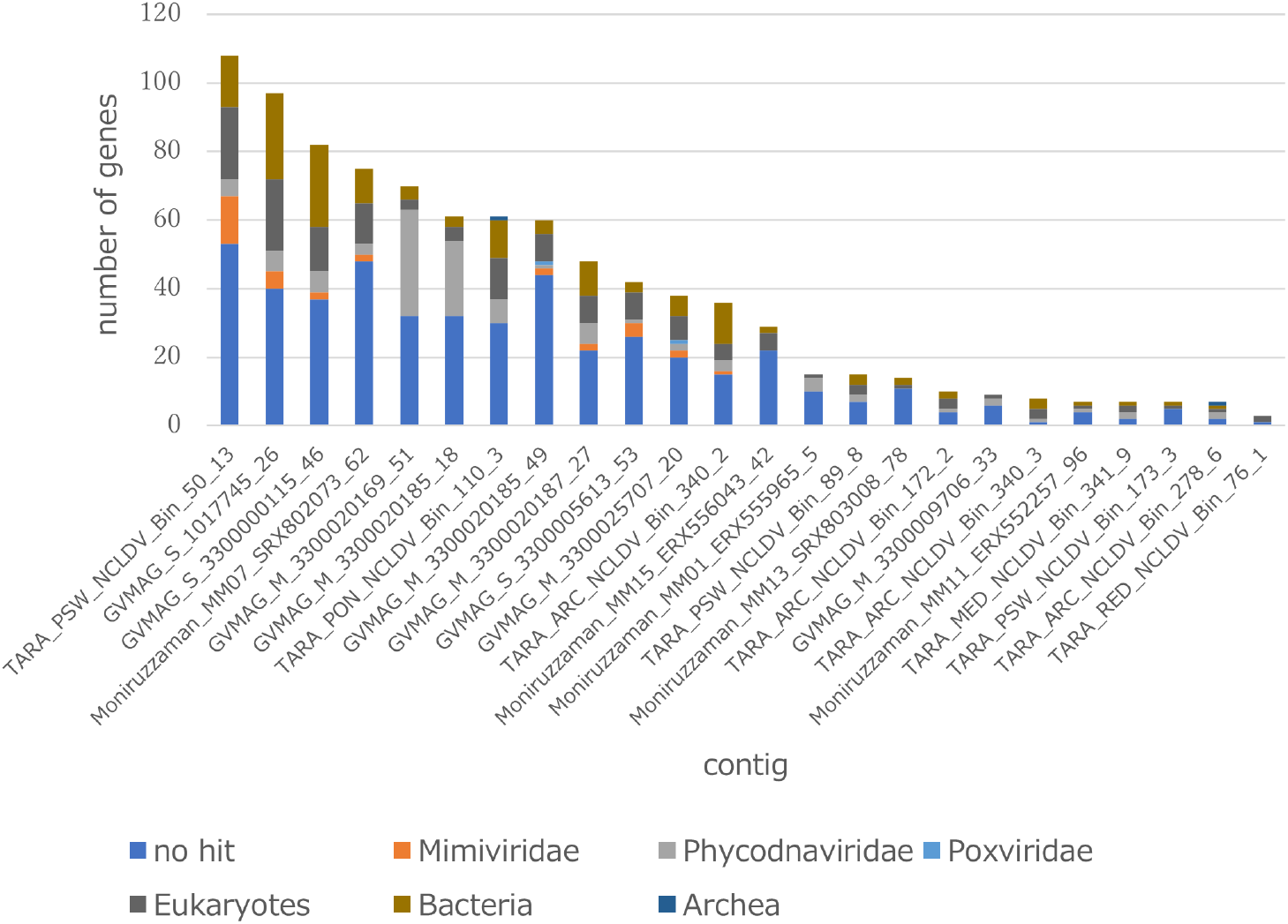
Taxonomic annotation of genes encoded in the virmyosin-harboring NCLDV contigs. Best hit-based taxonomic annotation was performed for each gene using BLAST against the RefSeq database.

To investigate the evolutionary relationships between the virmyosin-encoding NCLDVs, we performed phylogenetic analyses based on DNA polymerases (PolBs), a commonly used phylogenetic maker for NCLDVs. Eighteen of the 24 virmyosin-harboring MAGs were found to also encode PolB and were thus subjected to this analysis. The generated tree indicates that all but one of these virmyosin-encoding MAGs belong to the *Imitervirales* order, the grouping of which was supported with a bootstrap value of 91% (**Figure 2**). However, the lineages of virmyosin-encoding MAGs were scattered within at least four clades of the *Imitervirales* branches. The *Imitervirales* MAGs previously shown to encode viractins did not show close relationships with the virmyosin encoding MAGs. One of the virmyosin-encoding MAGs was identified within the *Phycodnaviridae* family.

**Figure 2.**
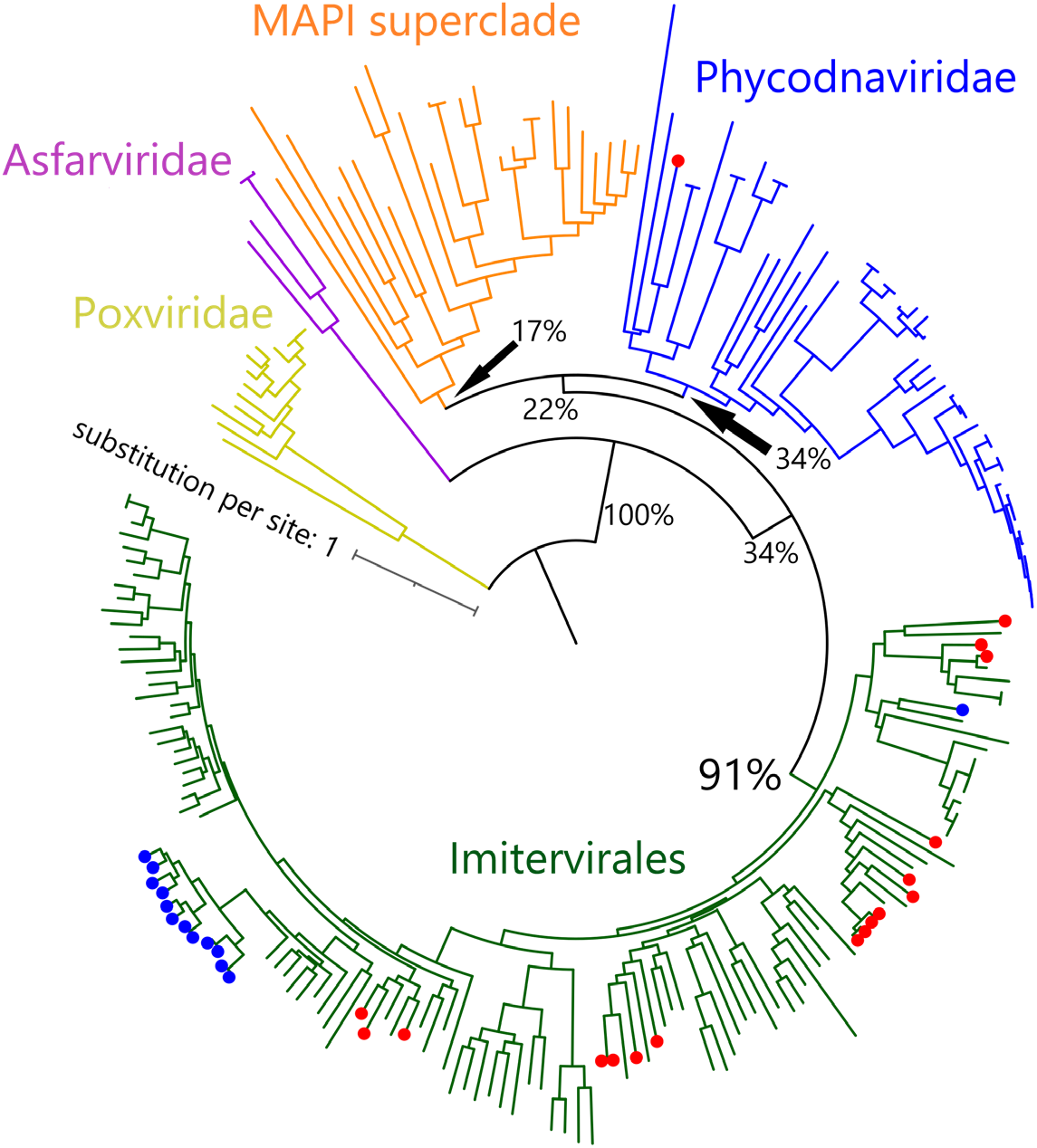
Phylogenetic tree of NCLDVs based on DNA polymerase B. The tree was built from 219 PolB sequences of NCLDVs (18 MAGs encoding myosin gene, 13 MAGs encoding actin gene, and 188 other broad and unbiased taxa from the reference database). Branches labeled with red and blue circles represent NCLDV MAGs with virmyosin and viractin genes, respectively. Branches are color-coded based on family- or order-level taxonomy. The MAPI superclade (shown in orange) comprises *Marseilleviridae, Ascoviridae*, Pitho-like viruses and *Iridoviridae*. The bootstrap value for the branch between *Imitervirales* and other families is 91%. We used *Poxviridae* as an outgroup to root the tree. The LG+F+R10 substitution model was selected by IQ-TREE for the best model for tree reconstruction.

Since myosins are universal in the eukaryotic domain, we hypothesized that these NCLDVs acquired myosin homologs by horizontal gene transfers (HGTs) from various eukaryotes. To determine the source eukaryotic lineages for the putative HGTs, we performed phylogenetic analyses on the virmyosin and reference myosin sequences. We generated four phylogenetic trees with different methods (**Figure 3, Figure S2**, see Materials and Methods). The phylogenetic placement of the virmyosin from the MAG belonging to the *Phycodnaviridae* family was unstable. However, all the trees showed a monophyletic group of *Imitervirales* virmyosins. The grouping is supported by a bootstrap value of 71% (**Figure S2a**).

**Figure 3.**
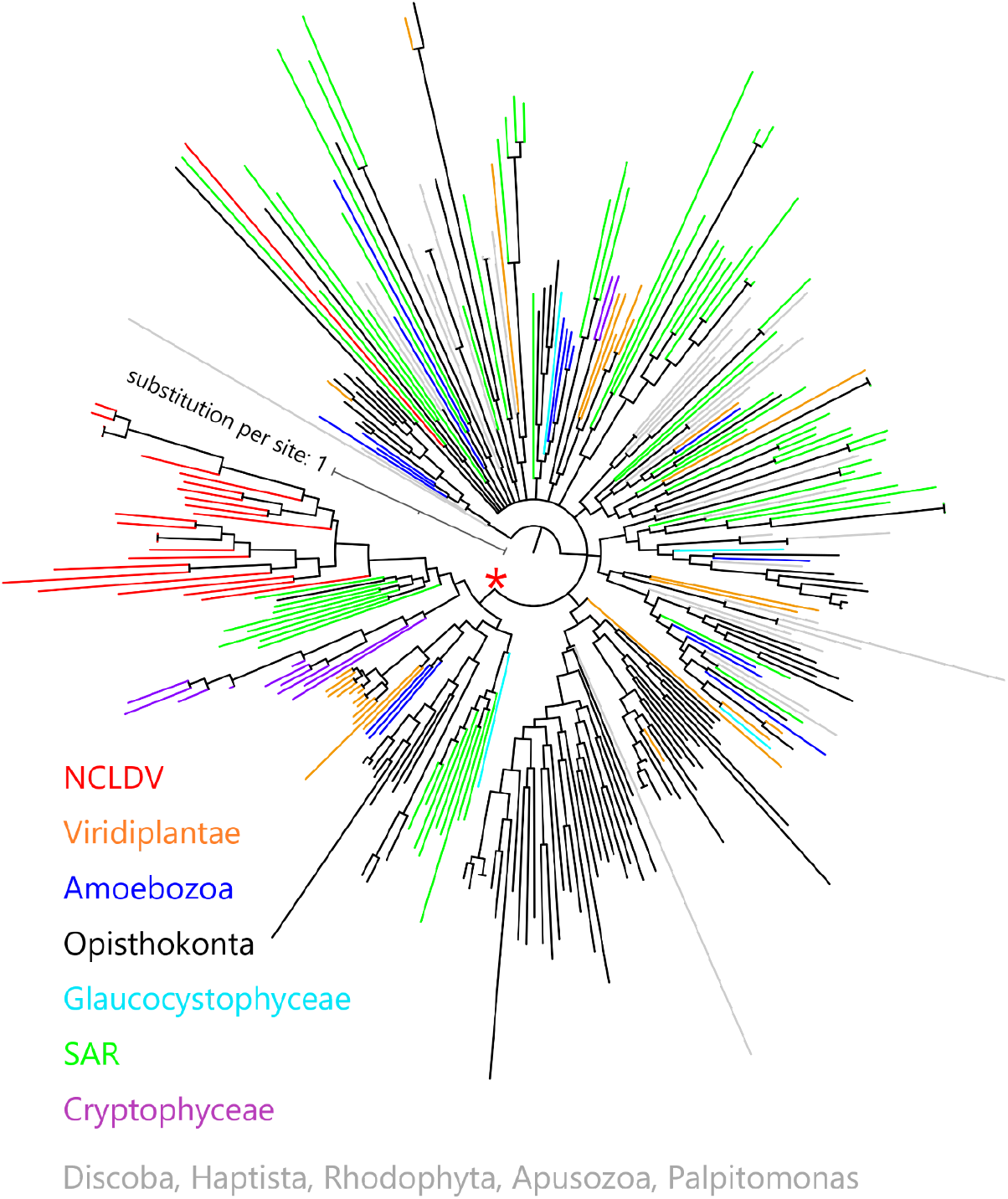
Phylogeny of myosin from NCLDVs and eukaryotic lineages. The tree was built from the multiple sequence alignment of 286 myosin head domain sequences. Branches are color-coded based on taxonomic groups (red branches correspond to virmyosins). We consider myosin-II as an outgroup to root this tree. The LG+R10 substitution model was selected by IQ-TREE for the best model for tree reconstruction.

To examine the effect of long-branch attraction on the formation of the *Imitervirales* virmyosin clade, we computed the pair-wise sequence identities within virmyosins and between virmyosins and eukaryotic myosins (**Figure 4**). The sequence similarities for many of the virmyosin pairs were higher than those between virmyosins and eukaryotic myosins, thus diminishing the possibility of long branch attraction effect on the monophyletic grouping the *Imitervirales* virmyosins. The *Imitervirales* virmyosin clade is branching out from orphan myosins of the SAR (Stramenopiles, Alveolates, and Rhizaria) supergroup, but the grouping of these virmyosins and the orphan myosins was not supported. Finally, to further improve the tree reconstruction, we built a phylogenetic tree using the virmyosin sequences as well as eukaryotic myosin sequences that formed a clade together with the virmyosins (the clade marked with ‘*’ in **Figure 3**). The newly generated tree again displayed the monophyletic grouping of the *Imitervirales* virmyosins (bootstrap value, 98%) and placed it within a clade of myosin sequences from Stramenopiles (the diatom *Thalassiosira pseudonana*, and the oomycete *Hyaloperonospora parasitica*) and a fungus (*Rhizopogon burlinghamii*) (**Figure 5, Figure S2e**). The grouping between the virmyosins and the eukaryotic myosins was not supported (bootstrap value, 15%), leaving the origin of the vimyosins unresolved.

**Figure 4.**
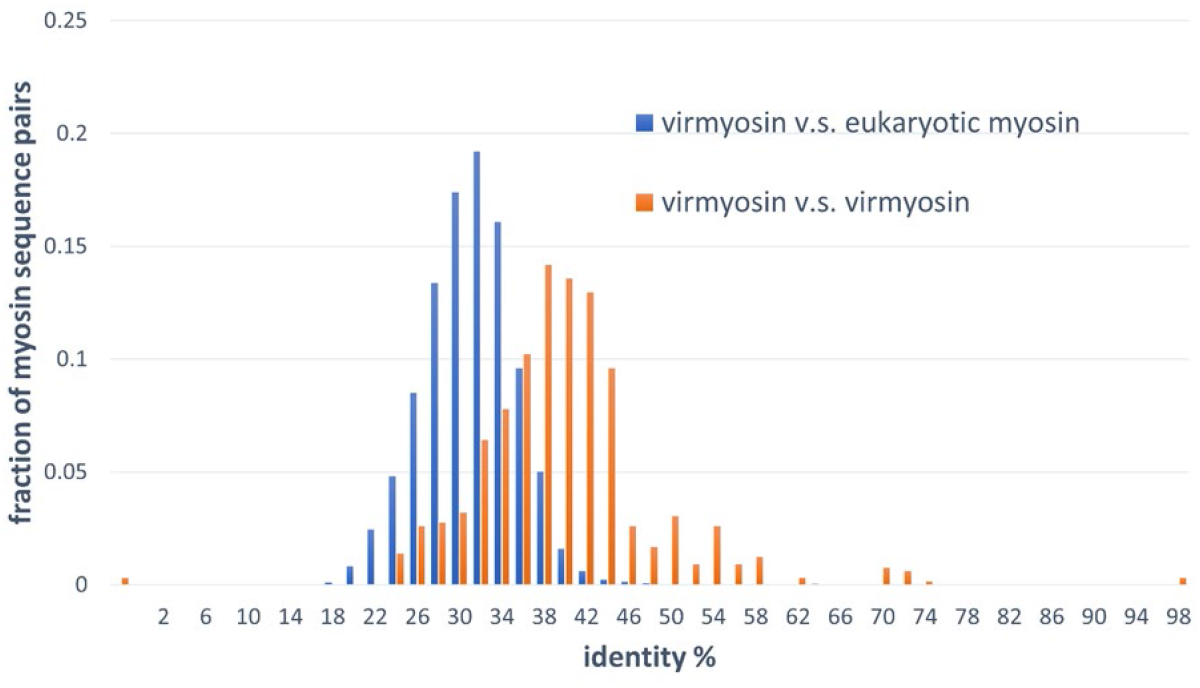
Distributions of pair-wise myosin sequence similarities within the NCLDVs (virmyosin *vs* virmyosin) and between NCLDVs and eukaryotes (virmyosin *vs* eukaryotic myosin). This graph shows that the mean value of myosin similarity within NCLDVs is higher than that between NCLDVs and eukaryotes.

**Figure 5.**
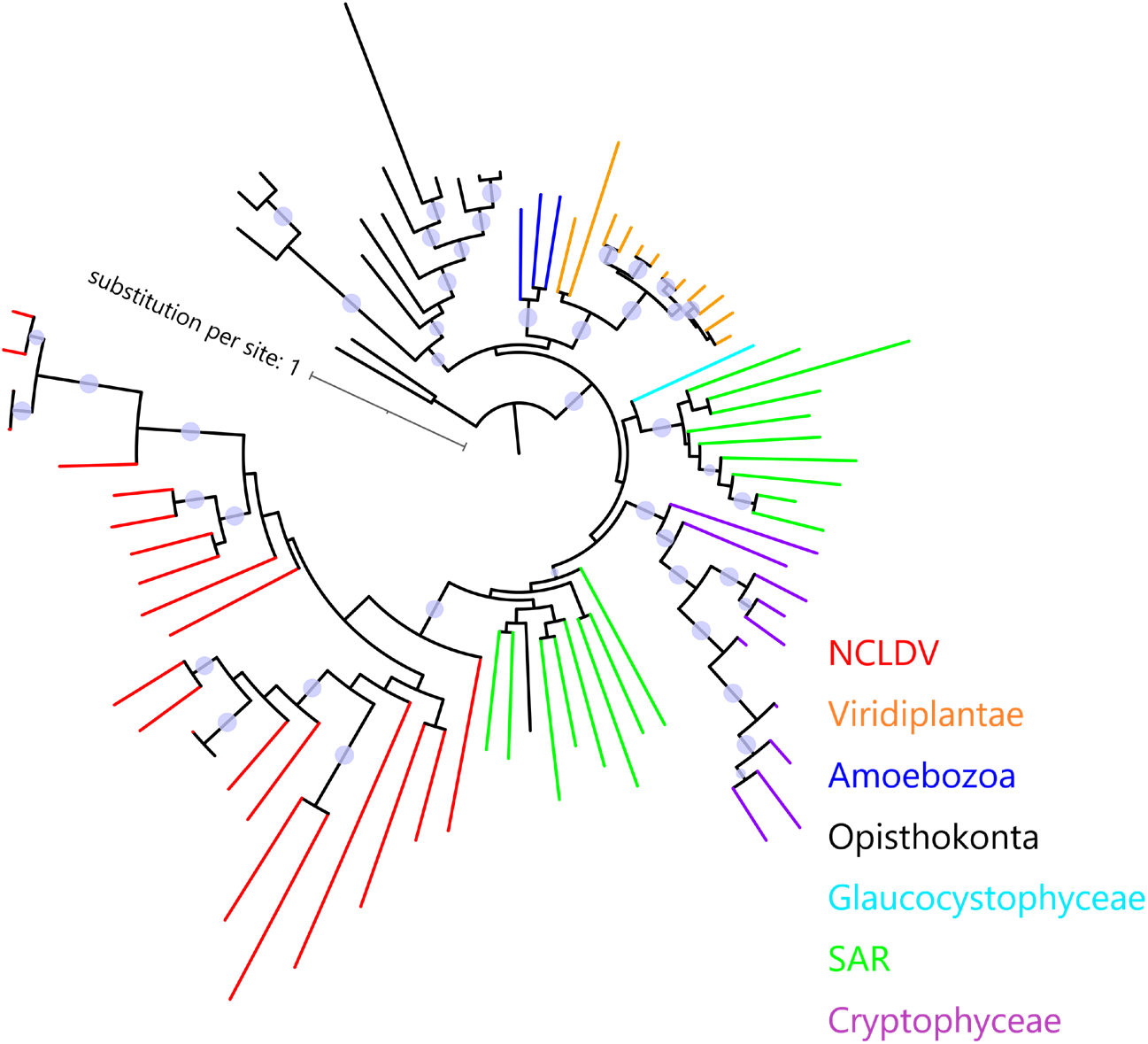
Phylogeny of virmyosins and their close relatives in eukaryotes. The tree was built from 81 myosin sequences of NCLDVs, Amoebozoa, SAR, Opisthokonta, Cryptophyceae, Viridiplantae and Glaucocystophyceae. Branches are color-coded based on taxonomic groups (red branches correspond to virmyosins). We consider myosin-II of *Nasonia vitripennis* and *Neurospora crassa* as outgroups to root this tree. Bootstrap values higher than 80% are represented as circles on the branches. The LG+F+R7 substitution model was selected by IQ-TREE for the best model for tree reconstruction.

### Incongruence of phylogenetic trees between myosins and DNA polymerases

We next examined the congruence of the virmyosin and PolB trees by focusing on the 18 virmyosins that were found in MAGs that encode PolB. Congruence of the topologies between these trees is expected if the original virmyosin gene was acquired by an ancestral virus prior to the divergence of viral clades or families. We generated a new phylogenetic tree based on the 18 virmyosin sequences and the 18 PolB sequences. To investigate the congruence of the trees, we focused on clades in the virmyosin tree that were supported with a bootstrap value greater than 90% (**Figure 6**). The analysis revealed that several groups of closely related sequences [i.e., (TARA_PSW_NCLDV_Bin_50_13, GVMAG_M_3300020185_49); (Moniruzzaman_MM07_SRX802073_62, GVMAG_S_1017745_26); (TARA_MED_NCLDV_Bin_341_9, GVMAG_M_3300020185_18); (GVMAG_S_3300000115_46, TARA_ARC_NCLDV_Bin_340_2, TARA_ARC_NCLDV_Bin_340_3, Moniruzzaman_MM13_SRX803008_78)] form monophyletic groups also in the PolB tree. However, these two trees were incongruent at deeper branches. For instance, one of the monophyletic clades in the virmyosin tree composed of four MAGs [i.e., (Moniruzzaman_MM07_SRX802073_62, GVMAG_S_1017745_26, GVMAG_M_3300020169_51, GVMAG_S_3300005613_53)] turned out to be polyphyletic with statistical supports in the PolB tree (**Figure 6**). This result suggests that multiple and distantly related ancestral viruses of *Imitervirales* independently acquired myosin genes through HGTs.

**Figure 6.**
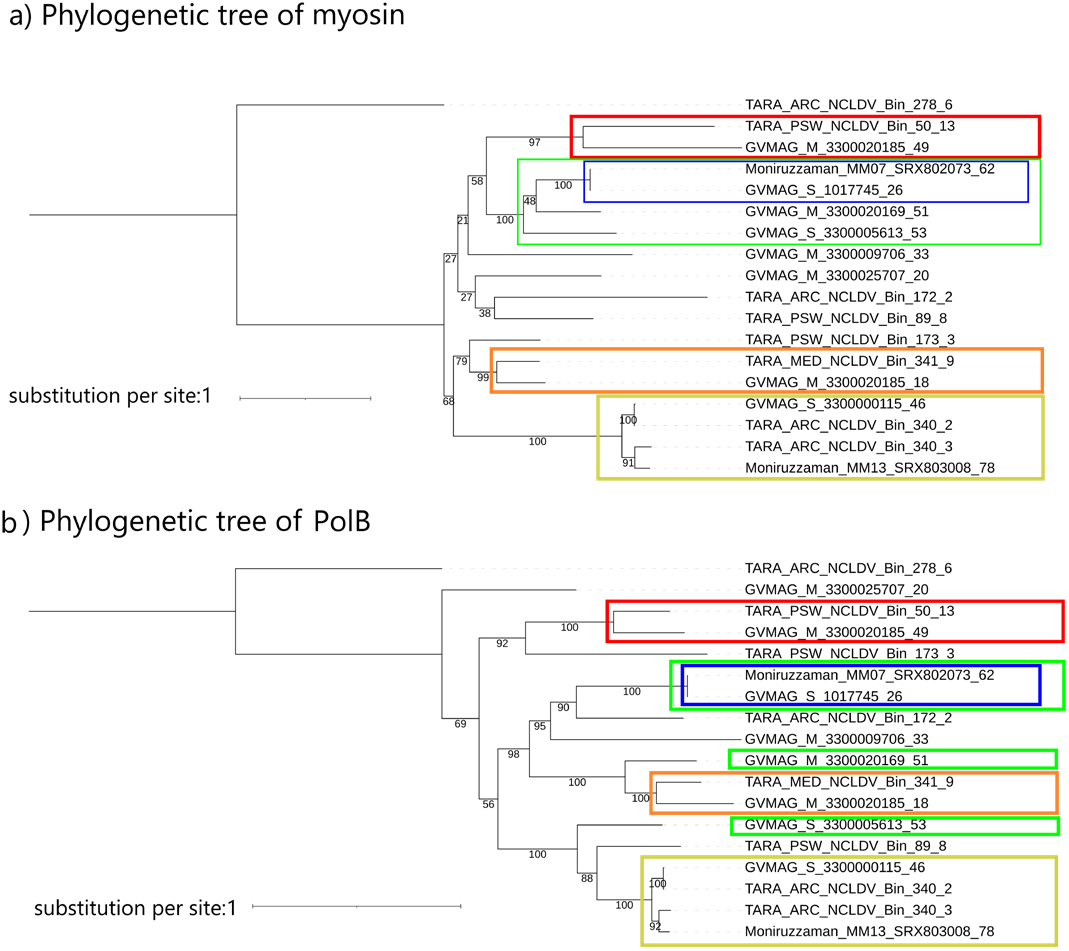
Phylogenetic trees of (a) myosin and (b) PolB sequences from MAGs encoding both genes. Labels at the leaves are the identifiers of the NCLDV MAGs. Clades supported with a bootstrap value >90% in the myosin tree are marked in both trees by colored rectangles. The LG+F+I+G4 substitution model was selected by IQ-TREE for the best model for both tree reconstructions.

The monophyletic grouping of the *Imitervirales* virmyosins and the scattered distribution of the virmyosin-encoding *Imitervirales* in the PolB tree are intriguing. We consider three major possibilities for the origin of virmyosins. The first possibility is that all the *Imitervirales* myosin genes were vertically inherited from an ancestral virus of the *Imitervirales*, and subsequently lost independently in most of the descendant lineages. This scenario is not parsimonious as it assumes many independent gene losses. The second possibility is that myosin genes were independently recruited multiple times from eukaryotes by ancestral viruses belonging to the *Imitervirales*. This scenario can account for the topological differences in the two trees. However, this scenario cannot readily explain the monophyletic grouping of the virmyosins, as this scenario needs to additionally assume independent horizontal acquisitions of myosin genes by distantly related ancestral viruses from the same or closely related eukaryotes (e.g., ancestral Stramenopiles). Such acquisitions seem implausible because host changes are likely rampant events for *Imitervirales* given the wide host ranges of the known viruses in this group (Sun et al., 2020). The third possibility is that a myosin gene transfer occurred once in a viral lineage of *Imitervirales* from its host. Then, after the viral myosin gene acquired beneficial functions for the virus, this gene was transferred to other viruses, probably during co-infection in the same host (which may be different from the current hosts). Of note, a previous study reported a clear case of horizontal gene transfer between NCLDVs (Christo-Foroux et al., 2020). This last scenario can explain both the topological difference between the virmyosin and PolB trees and the monophyletic grouping of the virmyosins. Further exploration of the actual diversity of *Imitervirales*, and more globally of NCLDVs, from various environments will certainly provide important insights regarding the robustness of these hypotheses

### Kinesin genes are associated with virmyosin genes

We identified 880 protein families/domains that showed a significant level of co-occurrence with virmyosin genes among the MAGs of the *Imitervirales* order by performing chi-square test (false discovery rate 0.01; **Table S2**). Interestingly, kinesins were significantly enriched in the *Imitervirales* MAGs encoding virmyosin (*q*-value = 2.9e-33, ranked at 7-th). Of 24 MAGs encoding virmyosins, 16 (67%) encoded kinesins, a score that might be underestimated due to the possible incomplete nature of NCLDV MAGs. Myosins are known to walk along actin filaments, which are located at the peripheric side of the cytoplasm, whilst kinesins walk toward the outer side of the cytoplasm along microtubules, which are mainly located in the inner side of the cytoplasm. Both kinesins and some classes of myosins function as a transporter to carry specific materials on these filaments. Viral kinesins may function to transport viral particles along microtubules and pass them to viral myosins at the end of microtubules. Myosins then transport viruses to the outer side of the cell to finally export the viral particles outside the cell. Intracellular enveloped viruses of vaccinia virus are transported towards the cell surface in a microtubule-dependent process (Hollinshead et al., 2001). It is possible that the viral myosins and kinesins found in NCLDV MAGs may function in a similar viral particle transport process.

## CONCLUSIONS

In this study, we provided strong evidence showing that marine members of NCLDVs encode myosin homologs (virmyosins), which contain the head domain but lack the tail domain. The function of virmyosin could not be inferred based on the similarity to functionally characterized myosin homologs. However, virmyosin genes were frequently accompanied by kinesin genes, another class of motor proteins widespread in the eukaryotic domain. Together with the previously discovered actin homologs in NCLDVs, these results suggest that the genetic independence of NCLDVs from their hosts encompasses a wide-range of cellular processes, including intracellular trafficking as implied by our study and the translation process as considered from previous discoveries of translation genes in this group of viruses. Our phylogenetic analyses suggest a complex evolutionary origin of the virmyosin genes, which may involve not only HGTs from eukaryotes to NCLDVs but also intra-virus HGTs within NCLDVs. The functions encoded in the huge genetic pool of NCLDVs are revealing the amazingly diverse strategies to control host cellular processes in this diverse group of viruses. Scrutinizing available and newly generated environmental genomes will contribute to better characterizing the infection strategies of this fascinating group of viruses.

## ACKNOWLEDGEMENTS

This study was supported by JSPS KAKANHI (18H02279) and Research Unit for Development of Global Sustainability, Kyoto University Research Coordination Alliance. Computation time was provided by the SuperComputer System, Institute for Chemical Research, Kyoto University.

