## Supplementary figure S1 for "Discovery of viral myosin genes with complex evolutionary history within plankton"

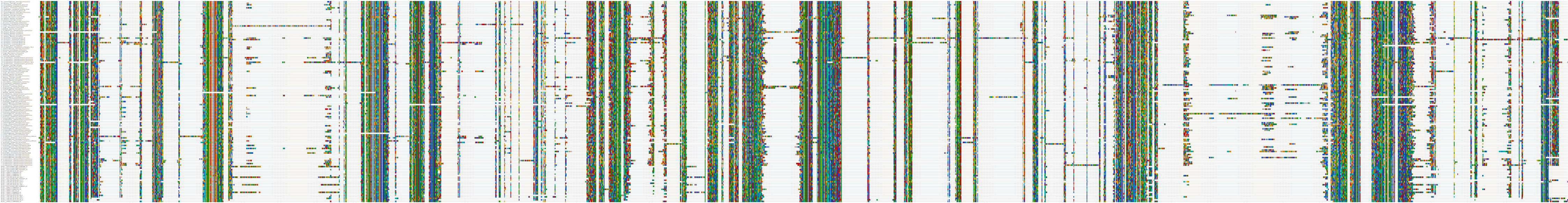

Supplementary Figure S1: homology of head domain of virmyosin and myosin sequences from various eukaryotes. We identified myosin head domain and create their multiple alignment.
