## Supplementary figure S2 for "Discovery of viral myosin genes with complex evolutionary history within plankton"

**Supplementary Figure 2.** Phylgenetic trees of myosin sequences. **(a)** Phylogenetic tree of myosins (gappy/IQ-TREE). The tree was built from the multiple sequence alignment of 286 myosins. Tree scale represents substitution per site. We consider myosin-II as outgroup to root this tree. **(b)** Phylogenetic tree of myosins (strict/IQ-TREE). The tree was built from the multiple sequence alignment of 286 myosins. Tree scale represents substitution per site. We consider myosin-II as outgroup to root this tree. **(c)** Phylogenetic tree of myosins (gappy/RAxML). The tree was built from the multiple sequence alignment of 286 myosins. Tree scale represents substitution per site. We consider myosin-II as outgroup to root this tree. **(d)** Phylogenetic tree of myosins (strict/RAxML). The tree was built from the multiple sequence alignment of 286 myosins. Tree scale represents substitution per site. We consider myosin-II as outgroup to root this tree. **(e)** Phylogenetic tree of myosins (gappy/IQ-TREE). The tree was built from the multiple sequence alignment of 82 myosins. Tree scale represents substitution per site. We consider myosin-II as outgroup to root this tree.

### Supplementary Figure 2a

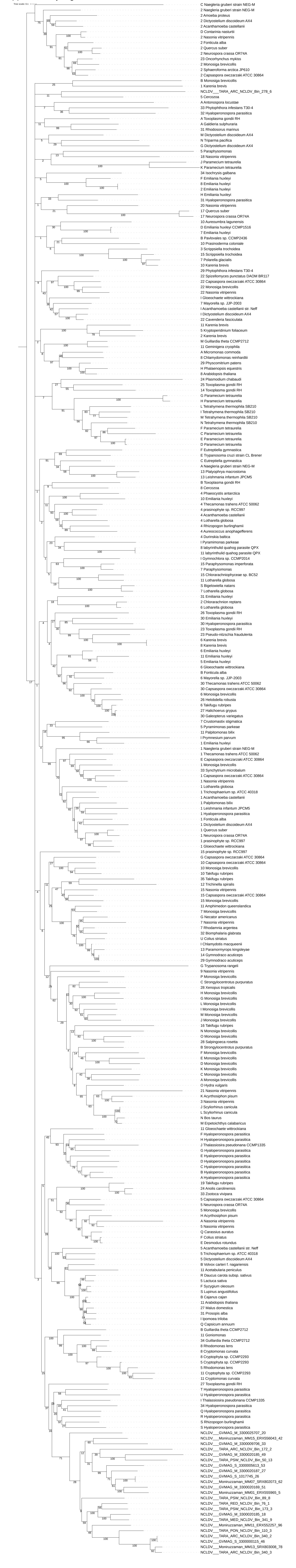

### Supplementary Figure 2b

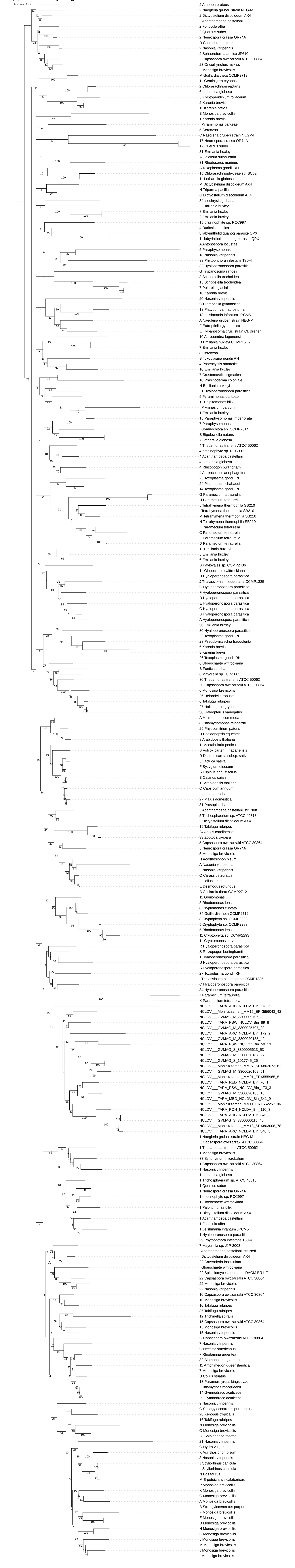

### Supplementary Figure 2c

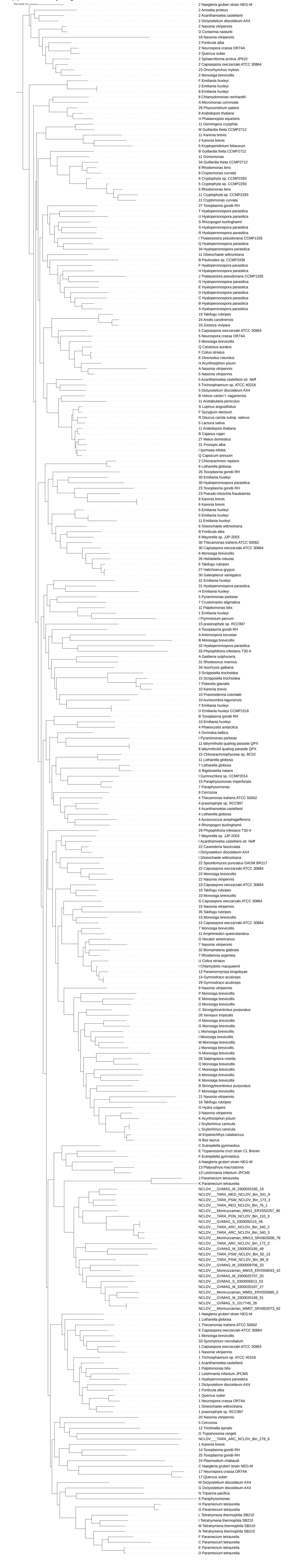

### Supplementary Figure 2d

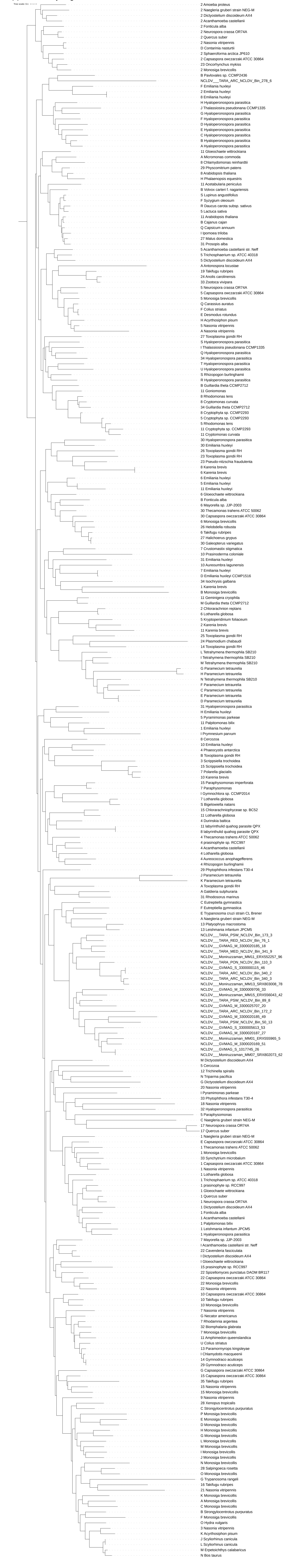

### Supplementary Figure 2e

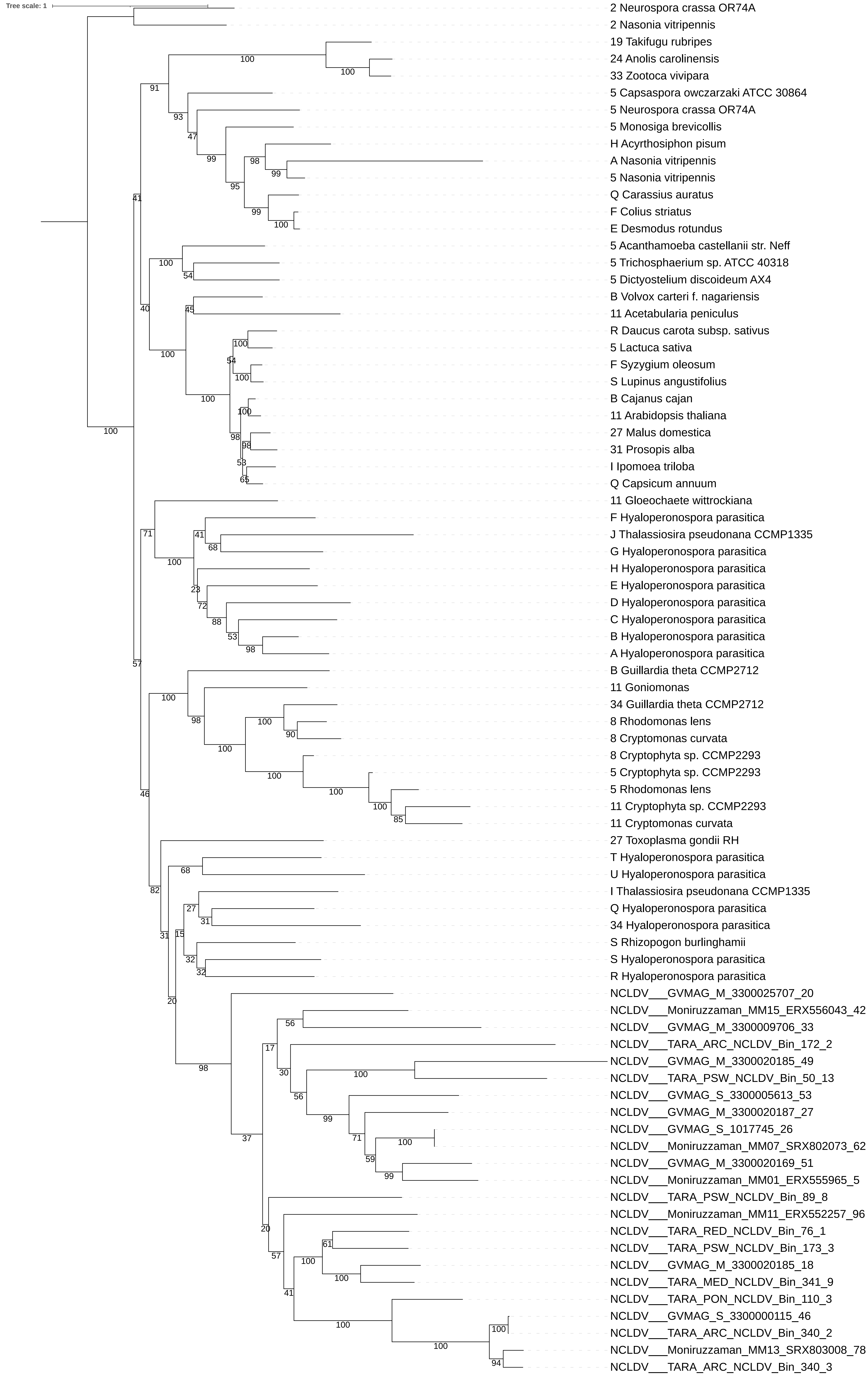
